# Development of an STR panel for a non-native population of an endangered species

**DOI:** 10.1101/2021.05.13.442066

**Authors:** Erin Hill, Adrian Linacre, Simon Toop, Nicholas Murphy, Jan Strugnell

## Abstract

The establishment of non-native populations of threatened and legally protected species can have many implications for the areas where these species have been introduced. Non-native populations of threatened species have the potential to be exploited and therefore the subject of legal protection, while conversely, if they have become invasive in their introduced range, there is the likelihood that population control will be carried out to reduce abundance and negative impacts associated with introduced species. From both a legal and invasive species monitoring standpoint, it is important to know how many individuals are present. Short tandem repeats (STRs) were developed for the hog deer, an endangered species that was introduced following European settlement to Victoria, Australia using Illumina MiSeq sequencing technology. These markers were combined with previous STRs characterised for hog deer to create a 29-plex identification system. A total of 224 samples were genotyped across the population in Victoria, and further analyses of null allele frequencies, deviation from Hardy-Weinberg equilibrium, and the removal of monomorphic or low amplifying markers resulted in a final marker panel of 17 loci. Despite low values for number of alleles at each locus (2-4), probability of identity showed sufficient discrimination power, with an average probability of identity at 9.46 × 10^−7^, and a probability of sibling identity of 7.67 × 10^−4^ across all sites. These findings show that it is feasible to create an informative DNA profiling system that can distinguish between individuals for applications in both wildlife forensic and population control research.

## Introduction

The establishment of non-native populations of threatened and legally protected species can have many implications for the areas where these species have been introduced [1]. Non-native populations of threatened species have the potential to be exploited and therefore the subject of legal protection, and may be targeted for trophies, meat, or as part of the pet or timber trade [2-4]. In Australia, a number of ungulate species were introduced by Acclimatisation Societies in the 19^th^ century are now threatened with extinction in their native range; rusa deer (*Rusa timorensis*) and sambar deer (*Rusa unicolor*) are listed as vulnerable by the IUCN, while banteng (*Bos javanicus*) and hog deer (*Axis porcinus*) are listed as endangered. Although these species are threatened with extinction in their native ranges, they are legal to hunt within their introduced ranges. These species are often exploited in their native range for meat and trophies, and these pressures can extend to the non-native populations as well.

Hog deer are CITES listed, endangered within their native range, but also considered an invasive species across Australia, and protected wildlife and a managed game species in Victoria. Hunting restrictions of hog deer are enforced in Victoria, with a limited one-month hunting season (April), bag limits of only one male and female deer per hunter each season, and additional tagging, inspection and reporting requirements [5]. Despite these strict regulations, illegal harvesting of hog deer occurs as trophies and meat are highly sought after by hunters both in Australia and overseas due to the rarity of the species. While hog deer are considered protected wildlife in Victoria, population control of the species can also take place, and has been occurring at Wilsons Promontory National Park since 2015 [6]. From both a legal and population control perspective, it is important to be able to distinguish between individuals. Game Officers often encounter illegally harvested deer in multiple body parts or stored with other species, which can make it difficult to determine how many individuals have been illegally taken (Toop pers. comm.). Conversely, invasive species managers may be interested in knowing how many individuals are present at a given location and monitor particular sites over time in order to observe changes in population sizes following population control. An effective DNA profiling system that is able to distinguish between individuals is therefore necessary to address many of the management issues surrounding hog deer in Victoria. DNA profiling in wildlife forensic cases typically uses short tandem repeats (STRs), which have been shown in the past to be useful for both individual assignment and population assignment [7-9] and can provide a fast turn-around of results that may not be possible when using other marker types, such as SNPs [10]. STRs can be highly polymorphic, and when multiple STRs are used they can provide high discriminatory power for identifying individuals. This study describes a set of polymorphic STRs developed for the hog deer population present in Victoria, Australia, and their suitability for distinguishing between individuals.

## Methods

A total of 231 hog deer tissue samples, including two voucher specimens from Museum Victoria (ref. no. Z52264 and Z52231), were collected across the hog deer distribution in the Gippsland region of Victoria by hunters between 2008-2017. Extractions were performed using a DNeasy Blood and Tissue Kit (Qiagen) following the manufacturer’s instructions, with negative controls run throughout. DNA quantity was measured using Qubit 2.0 Fluorometer (Invitrogen).

STR regions were identified using an Illumina *de novo* sequencing approach. Paired-end libraries were prepared for four hog deer samples collected from Sunday Island in 2015 using a NebNext Ultra Library Prep Kit (BioLabs), with size selection of 500-600 base pairs (bp). Sequencing was performed on an Illumina MiSeq using a V3 (2×300) MiSeq Reagent Kit (Illumina). An average of 7 million reads from each sample were obtained from the sequencing run. Sequences were processed using the program *Kraken v0*.*10*.*5* [11] to remove non-target DNA sequences, resulting in an average of 5 million reads per sample. Samples were then processed in two ways to identify STRs at a range of base pair (bp) lengths. Firstly, the sample that contained the highest number of reads (∼9 million reads) was trimmed to remove low quality reads in *CLC Genomics Workbench 7* (CLC bio, Inc.) and then run through the program *Msatcommander 1*.*0*.*8* [12] to identify STRs and create primers. The second method involved *de novo* assembly of all four samples, using the program *CLC Genomics Workbench 7* (CLC bio, Inc.), with a minimum contig size of 1000. The consensus assembly was then used in the program *Msatcommander 1*.*0*.*8*. These two methods were implemented to identify STRs at a range of sizes, to be used later for multiplexing. *Msatcommander 1*.*0*.*8* was run to identify only tri-, tetra-, penta- and hexa-repeat units, as STRs of these repeat lengths are less prone to allele dropout and stuttering [13]. A pool of 725 potential STRs were identified using the first method, while 109 were identified through *de novo* assembly.

Primer pairs identified by *Msatcommander 1*.*0*.*8* were then tested for successful amplification, and those that successfully amplified a DNA fragment were then tested for polymorphism using 20 hog deer tissue samples across the species distribution in Victoria, including the two museum voucher specimens. PCR was performed in a total volume of 12.5 μL, consisting of 6.25 μL of MyTaq Red Mix (Bioline), 0.075 μL of forward primer (10 μM), 0.25 μL of reverse primer (10 μM), 0.085 μL of fluorescently labelled dye (6-FAM, VIC, or PET), 4.84 μL of H_2_O, and 1 μL of template DNA, ranging from 1.1 - 26.7 ng/μL. Cycling conditions were 95°C for 5 minutes, 35 cycles of 95°C for 30 seconds, annealing step for 30 seconds, 72°C for 1 minute, followed by a final extension of 72°C for 10 minutes. Annealing temperature varied depending on the primer pairs. Negative controls were run throughout the PCR process. Samples were sent to the Australian Genome Research Facility (AGRF) for genotyping and were visualised using *Geneious 9*.*0*.*5* [14] to determine polymorphism.

To reduce laboratory processing time, polymorphic STRs identified using the above method, as well as previously developed hog deer STRs that successfully amplified from [15] were pooled into three multiplexes. All primers were run through the program *AutoDimer* [16] to identify any potential primer dimer interactions between primers. Fluorescently tagged primers were then prepared to stock solutions according to the Qiagen Multiplex PCR Kit manual. PCRs were then performed in multiplexes with an initial primer concentration of 0.2 μM. PCR was carried out in 25 μL reactions, containing 12.5 μL of Qiagen Multiplex PCR master mix (Qiagen), 3.5 μL of 2 μM primer mix, 3.5 μL of Q-solution (Qiagen), 1.5 μL of RNase-free water (Qiagen), and 4 μL of 1-2 ng/μL template DNA. Cycling conditions were 95°C for 15 minutes, 35 cycles of 94°C for 30 seconds, annealing step for 90 seconds, 72°C for 2 minutes, followed by a final extension of 60°C for 30 minutes. Annealing temperature varied between 58°C - 61°C depending on the multiplex (Table 1). Primer concentrations were adjusted by increments of 0.1 μM for primer pairs that produced too strong, or weak final products (Table 1). Once primer concentrations were established for each multiplex, all 231 hog deer samples were diluted to 1-2 ng/μL and genotyped using the multiplex protocol described above, with positive and negative controls run throughout and sent to AGRF for genotyping.

**Table 1.**
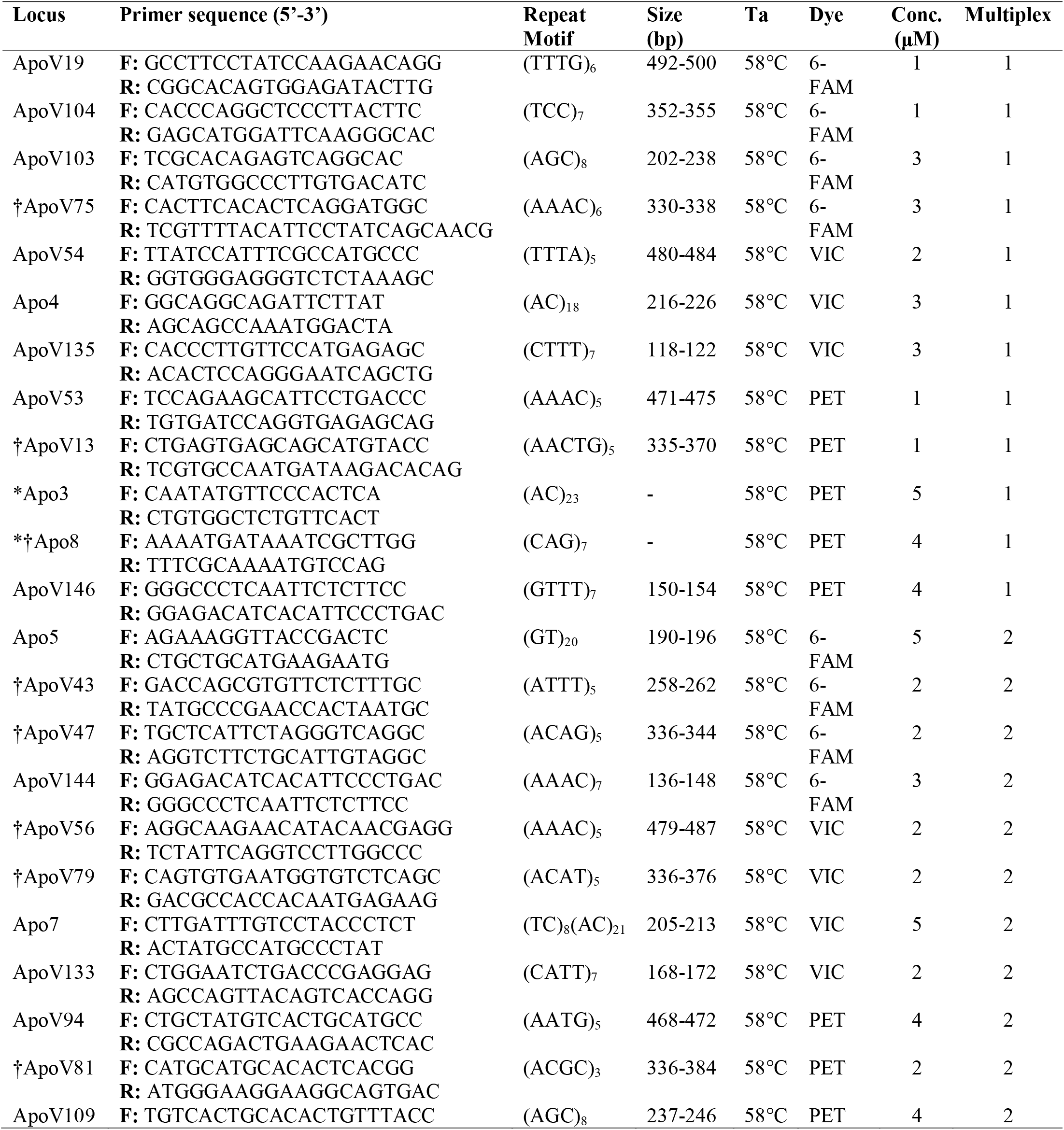
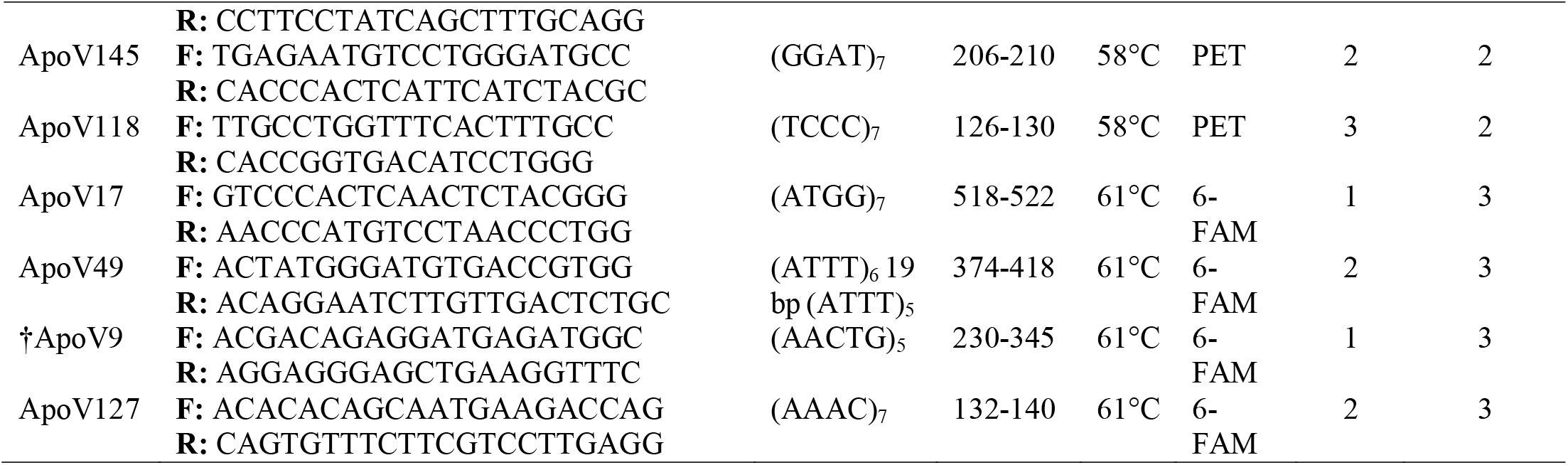
Characterisation and multiplex design of 24 newly developed STR markers for the hog deer, and five previously published hog deer STRs (Apo4, Apo3, Apo8, Apo5, and Apo7; [15]). All markers were tested on an initial 20 samples to confirm polymorphism. Loci with an * symbol were excluded from further analysis due to low amplification success; Loci with an † symbol were excluded from further analysis as they were found to be either monomorphic or produced more than two peaks in hog deer

Genotypes were visualised and scored using *Geneious 9*.*0*.*5* [14]. Samples that contained missing data at seven or more loci were removed from the dataset, leaving 224 samples for further analysis. Each STR was tested for Hardy-Weinberg equilibrium and linkage disequilibrium in the program *Genepop 4*.*2* [17]. The program *FreeNa* [18] was used to calculate the frequency of null alleles for each STR. The number of alleles and observed and expected heterozygosity was determined for each STR using the program *GenAlEx 6*.*502* [19], while polymorphism information content (PIC) was calculated using *Cervus 3*.*0*.*7* [20]. *GDA 1*.*0* was used to calculate F_IS_ and overall θ [21], with these values used to estimate the average PI in *API-Calc 1*.*0* [22]. PI and PI_sibs_ were additionally calculated in *GenAlEx 6*.*502* [19].

## Results and Discussion

A total of 142 STRs were tested for amplification success and polymorphism, and from these 24 polymorphic STRs were discovered and incorporated into three multiplexes, which also included five previously published hog deer STRs [15] (Table 1). Upon genotyping of all hog deer samples used in this study, a number of STRs included in the multiplexes were discovered to be unsuitable for further analysis. Previously published STRs Apo3 and Apo8 were discarded due to low amplification success, and samples that could be amplified at the Apo8 marker consistently amplified more than two peaks. This issue was also observed in ApoV43, where three peaks consistently amplified at this marker. Initial results testing polymorphism on 20 samples suggested that all markers described in this study were polymorphic in hog deer, however upon genotyping all available hog deer samples, a number of STRs were found to be monomorphic (ApoV9, ApoV13, ApoV47, ApoV56, ApoV75, ApoV79, and ApoV81), and it is likely that large stutter peaks contributed to these markers being misidentified as polymorphic.

Genotypes were successfully generated for 224 samples of hog deer, using 19 loci from the multiplexes described above. The number of alleles per locus (N_A_) ranged from 2-4, while polymorphism information content (PIC) varied from 0.01-0.64, with the lowest value attributed to ApoV17 and the highest to ApoV19 (Table 2). Low values of N_A_ have been described in hog deer STRs previously [15], likely due to these markers being developed and tested on a small, captive population of hog deer in China comprising approximately 30 individuals. It is worth noting however, that the present study observed a greater number of alleles than reported in [15] for the three markers that were successfully amplified across the dataset (Apo4, Apo5, and Apo7). Two loci were found to be homozygotes for different alleles and so comprised an observed heterozygosity of zero (ApoV17 and ApoV49). Excluding these loci, observed and expected heterozygosity ranged from 0.10-0.57 and 0.18-0.70 respectively.

**Table 2.** Number of alleles (N_A_), observed and expected heterozygosity (H_O_ and H_E_), polymorphism information content (PIC), Hardy-Weinberg Equilibrium (HWE), and frequency of null alleles (F_NULL_) for hog deer. Significant deviations from Hardy-Weinberg equilibrium are shown in bold; null allele frequencies above 5% are denoted with an asterisk (*). Loci with a † symbol were subsequently removed from further analysis

| Locus | n | $N_A$ | $H_O$ | $H_E$ | PIC | HWE | $F_{NULL}$ |
| --- | --- | --- | --- | --- | --- | --- | --- |
| ApoV19 | 224 | 4 | 0.57 | 0.70 | 0.64 | 0.33 | 0.079* |
| ApoV104 | 224 | 3 | 0.41 | 0.55 | 0.45 | 0.58 | 0.109* |
| ApoV103 | 224 | 2 | 0.47 | 0.37 | 0.30 | 0.12 | 0.000 |
| ApoV54 | 224 | 3 | 0.45 | 0.53 | 0.43 | 0.99 | 0.047 |
| Apo4 | 224 | 4 | 0.56 | 0.55 | 0.45 | 0.73 | 0.030 |
| ApoV135 | 224 | 3 | 0.36 | 0.42 | 0.33 | 0.91 | 0.047 |
| ApoV53 | 222 | 2 | 0.23 | 0.21 | 0.19 | 1 | 0.000 |
| †ApoV146 | 208 | 3 | 0.13 | 0.53 | 0.42 | **0.00** | 0.261* |
| Apo5 | 224 | 4 | 0.42 | 0.44 | 0.35 | 0.99 | 0.015 |
| †ApoV144 | 224 | 2 | 0.10 | 0.48 | 0.37 | **0.00** | 0.258* |
| Apo7 | 223 | 3 | 0.42 | 0.47 | 0.36 | 0.71 | 0.039 |
| ApoV133 | 224 | 2 | 0.39 | 0.40 | 0.32 | 0.96 | 0.008 |
| ApoV94 | 222 | 2 | 0.14 | 0.18 | 0.16 | 0.81 | 0.054* |
| ApoV109 | 224 | 3 | 0.21 | 0.20 | 0.19 | 1 | 0.000 |
| ApoV145 | 224 | 2 | 0.38 | 0.46 | 0.35 | 1 | 0.056* |
| ApoV118 | 224 | 2 | 0.35 | 0.41 | 0.33 | 0.76 | 0.050 |
| ApoV17 | 170 | 2 | 0.00 | 0.01 | 0.01 | 0.11 | 0.053* |
| ApoV49 | 213 | 3 | 0.00 | 0.14 | 0.13 | **0.00** | 0.181* |
| ApoV127 | 219 | 4 | 0.63 | 0.65 | 0.58 | 0.63 | 0.009 |

Significant linkage disequilibrium was detected in three pairwise comparisons between loci: ApoV144 and ApoV146 (Chi^2^ = >155.23, d.f. = 30, p = <7.87e-19), ApoV54 and ApoV53 (Chi^2^ = 49.06, d.f. = 24, p = 0.002), and ApoV104 and Apo5 (Chi^2^ = 52.08, d.f. = 30, p = 0.007). Significant deviations from Hardy-Weinberg equilibrium were detected in ApoV146, ApoV144, and ApoV49 (Table 2). Null alleles above 5% were detected in eight loci: ApoV19, ApoV104, ApoV146, ApoV144, ApoV94, ApoV145, ApoV17, and ApoV49. Given the evidence of significant linkage disequilibrium, significant deviations from Hardy-Weinberg equilibrium, and a high percentage of null alleles present in the STRs ApoV144 and ApoV146, these two markers were removed from further analyses of probability of identity. A number of loci across the three multiplexes described have been removed from further analysis owing to issues in amplification, monomorphic loci, and significant deviations from statistical tests. In order to further streamline lab processing of samples and overall cost, removal of uninformative STR loci form the multiplexes would be beneficial, which may additionally decrease the number of multiplexes needed for genetic analysis.

Seventeen loci were retained for probability of identity (PI) analyses of hog deer. The overall θ was 0.09, while the overall F_IS_ was 0.05; these values were used to calculate the average PI in *API-Calc 1*.*0*. The most and least informative loci for PI analysis varied based on the type of PI calculation being performed (PI or PI_sibs_ calculated in *GenAlEx 6*.*502*, and average PI calculated in *API-Calc 1*.*0*). However, STRs ApoV19 and ApoV127 were consistently identified as being in the top three most informative markers. The two least informative markers appear to be ApoV49 and ApoV17, however these STRs were not always identified as the least informative across all populations and calculations. PI and average PI were 4.44 × 10^−7^ and 9.46 × 10^−7^ respectively, while PI_sibs_ generated a final accumulative value of 7.67 × 10^−4^ (Fig.1). After the addition of nine loci, PI and average PI values were below the recommended threshold outlined by [23] for wildlife forensic cases across all populations, suggesting appropriate levels of discrimination between samples for forensic purposes. The total accumulative PI_sibs_ value was also within the range of this threshold and is comparable to other non-human forensic studies; analyses of European brown bears (*Ursus arctos*) revealed an overall PI_sibs_ value of 1.3 × 10^−4^, while evaluation of *Cannabis sativa* seized in Australia measured a PI_sibs_ value of 5.5 × 10^−4^ [24-25]. These results demonstrate that the STR assay developed in this study can distinguish between hog deer individuals, providing a useful method for genetic screening of hog deer individuals for both forensic and population control purposes.

**Fig.1.**
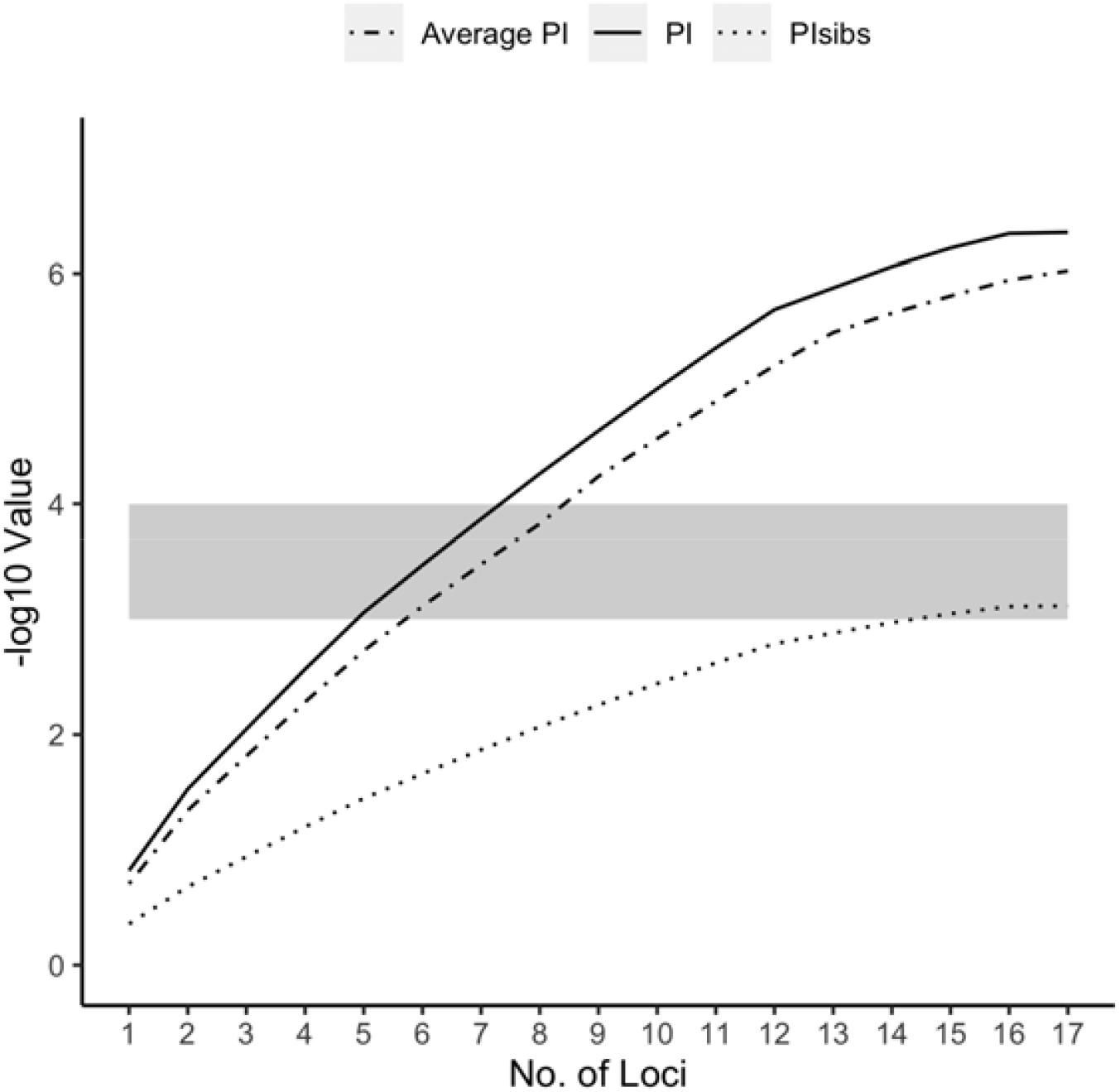
Hog deer accumulative PI values for based on 17 loci across all samples (n=224). Shaded grey area shows the threshold recommended by [23] to be informative for wildlife forensic applications. PI values have been - log10 transformed, so larger values represent a lower total PI score

## Acknowledgements

This project was jointly funded by the RFA grant ‘Securing Food, Water and the Environment’ (La Trobe University), and the Victorian Game Management Authority. We would like to thank Victorian Hog Deer Checking Station operators, Parks Victoria, Museums Victoria, Matt Salmon, Tim Thomas, and hunters for providing the hog deer samples used in this study. We also extend our thanks to Jude Hatley for providing assistance in operating the Illumina MiSeq.

## References

[1] Gibson, L., & Yong, D. L. (2017). Saving two birds with one stone: solving the quandary of introduced, threatened species. Frontiers in Ecology and the Environment, 15(1), 35–41

[2] Hall, R. J., Milner□Gulland, E., & Courchamp, F. (2008). Endangering the endangered: the effects of perceived rarity on species exploitation. Conservation Letters, 1(2), 75–81

[3] Alacs, E. A., Georges, A., FitzSimmons, N. N., & Robertson, J. (2010). DNA detective: a review of molecular approaches to wildlife forensics. Forensic Sci Med Pathol, 6(3), 180–194

[4] Kurland, J., Pires, S. F., McFann, S. C., & Moreto, W. D. (2017). Wildlife crime: a conceptual integration, literature review, and methodological critique. Crime Science, 6(1), 4

[5] Scroggie, M. P., Forsyth, D. M., & Brumley, A. R. (2012). Analyses of Victorian hog deer (Axis porcinus) checking station data: demographics, body condition and time of harvest. Arthur Rylah Institute of Environmental Research Technical Report Series, No. 230

[6] Game Management Authority (2017). Wilsons Promontory National Park Hog Deer Control Program. Game Management Authority. Victoria, Australia

[7] Wasser, S. K., Shedlock, A. M., Comstock, K., Ostrander, E. A., Mutayoba, B., & Stephens, M. (2004). Assigning African elephant DNA to geographic region of origin: applications to the ivory trade. Proc Natl Acad Sci U S A, 101(41), 14847–14852

[8] Frantz, A. C., Pourtois, J. T., Heuertz, M., Schley, L., Flamand, M. C., Krier, A., … Burke, T. (2006). Genetic structure and assignment tests demonstrate illegal translocation of red deer (Cervus elaphus) into a continuous population. Mol Ecol, 15(11), 3191–3203

[9] Hekkala, E. R., Amato, G., DeSalle, R., & Blum, M. J. (2009). Molecular assessment of population differentiation and individual assignment potential of Nile crocodile (Crocodylus niloticus) populations. Conservation Genetics, 11(4), 1435–1443

[10] Ogden, R. (2011). Unlocking the potential of genomic technologies for wildlife forensics. Mol Ecol Resour, 11, 109–116

[11] Wood, D. E., & Salzberg, S. L. (2014). Kraken: ultrafast metagenomic sequence classification using exact alignments. Genome Biology, 15(R46)

[12] Faircloth, B. C. (2008). msatcommander: detection of microsatellite repeat arrays and automated, locus-specific primer design. Mol Ecol Resour, 8(1), 92–94

[13] Linacre, A., Gusmao, L., Hecht, W., Hellmann, A. P., Mayr, W. R., Parson, W., … Morling, N. (2011). ISFG: recommendations regarding the use of non-human (animal) DNA in forensic genetic investigations. Forensic Sci Int Genet, 5(5), 501–505

[14] Kearse, M., Moir, R., Wilson, A., Stones-Havas, S., Cheung, M., Sturrock, S., … Drummond, A. (2012). Geneious Basic: an integrated and extendable desktop software platform for the organization and analysis of sequence data. Bioinformatics, 28(12), 1647–1649

[15] Lian, H., Yu, J.-Q., Ge, Y.-F., & Fang, S.-G. (2008). Nine novel microsatellite markers for the hog deer (Axis porcinus). Conservation Genetics, 10(3), 681–683

[16] Vallone, P. M., & Butler, J. M. (2004). AutoDimer: a screening tool for primer-dimer and hairpin structures. Biotechniques, 37(2), 226–231

[17] Raymond, M., & Rousset, F. (1995). GENEPOP (version 1.2): population genetics software for exact tests and ecumenicism. Heredity, 86, 248–249

[18] Chapuis, M.-P., & Estoup, A. (2006). Microsatellite null alleles and estimation of population differentiation. Molecular Biology and Evolution, 24(3), 621–631

[19] Peakall, R., & Smouse, P. E. (2012). GenAlEx 6.5: genetic analysis in Excel. Population genetic software for teaching and research--an update. Bioinformatics, 28(19), 2537–2539

[20] Marshall, T., Slate, J., Kruuk, L., & Pemberton, J. (1998). Statistical confidence for likelihoodLbased paternity inference in natural populations. Mol Ecol, 7(5), 639–655

[21] Lewis, P., & Zaykin, D. (2001). Genetic Data Analysis (GDA: version 1.0 d16c): A computer program for the analysis of allelic data. Website http://www.eeb.uconn.edu/people/plewis/software.php

[22] Ayres, K., & Overall, A. (2004). API-CALC 1.0: a computer program for calculating the average probability of identity allowing for substructure, inbreeding, and the presence of close relatives. Molecular Ecology Notes, 4, 315–318

[23] Waits, L. P., Luikart, G., & Taberlet, P. (2001). Estimating the probability of identity among genotypes in natural populations: cautions and guidelines. Mol Ecol, 10(1), 249–256

[24] Howard, C., Gilmore, S., Robertson, J., & Peakall, R. (2009). A Cannabis sativa STR genotype database for Australian seizures: forensic applications and limitations. Journal of forensic sciences, 54(3), 556–563

[25] Andreassen, R., Schregel, J., Kopatz, A., Tobiassen, C., Knappskog, P. M., Hagen, S. B., … Eiken, H. G. (2012). A forensic DNA profiling system for Northern European brown bears (Ursus arctos). Forensic Sci Int Genet, 6(6), 798–809

